# Biological age is increased by stress and restored upon recovery

**DOI:** 10.1101/2022.05.04.490686

**Authors:** Jesse R. Poganik, Bohan Zhang, Gurpreet S. Baht, Csaba Kerepesi, Sun Hee Yim, Ake T. Lu, Amin Haghani, Tong Gong, Anna M. Hedman, Ellika Andolf, Göran Pershagen, Catarina Almqvist, James P. White, Steve Horvath, Vadim N. Gladyshev

## Abstract

Aging is classically conceptualized as an ever-increasing trajectory of damage accumulation and loss of function, leading to increases in morbidity and mortality. However, recent *in vitro* studies have raised the possibility of age reversal. Here, we report that biological age is fluid and exhibits rapid changes in both directions. By applying advanced epigenetic aging clocks, we find that the biological age of young mice is increased by heterochronic parabiosis and restored following surgical detachment of animals. We also identify transient changes in biological age during major surgery, pregnancy, and severe COVID-19 in humans and/or mice. Together, these data show that biological age undergoes a rapid increase in response to diverse forms of stress, which is reversed following recovery from stress. Our study uncovers a new layer of aging dynamics that should be considered in future studies. Elevation of biological age by stress may be a quantifiable and actionable target for future interventions.

## Main Text

Biological age of organisms is thought to steadily increase over the life course. However, it is now clear that biological age is not indelibly linked to chronological age: individuals can be biologically older or younger than their chronological age implies.^1^ Moreover, increasing evidence in animal models and humans indicates that biological age can be influenced by disease,^2^ drug treatment,^3^ lifestyle changes,^4^ and environmental exposures,^5^ among other factors. Despite the widespread acknowledgment that biological age is at least somewhat malleable, the extent to which biological age undergoes reversible changes throughout life, and the events that trigger such changes, remain unknown.

DNA methylation (DNAm) clocks have emerged as the premier tool with which to assess biological age and begin to answer these questions. Such epigenetic aging clocks were innovated based on the observation that methylation levels of various subsets of CpG sites throughout the genome predictably change over the course of chronological age. First generation human DNAm clocks^6–8^ are constructed using machine learning approaches to build models trained on and designed to predict chronological age. Since the advent of DNAm clocks, both a suite of mouse DNAm clocks^3,9–13^ and second generation human DNAm clocks^14,15^ have emerged. Second generation human DNAm clocks integrate numerous phenotypic measures of aging (and, in some instances, chronological age) to produce a measure of morbidity/mortality risk and biological age. Another recently reported second generation approach, called DunedinPACE, uses longitudinal phenotypic training data to produce a measure of the rate of biological aging.^16,17^ DNAm clocks have excellent predictive ability and are responsive to known anti-aging/lifespan extending interventions such as caloric restriction.^11^ Although mechanistic questions on the nature of DNAm clocks remain, these clocks represent the current gold standard aging biomarker and are now widely utilized in the aging field, including in human clinical trials.^**18**^ Here, we leverage the power of DNAm clocks in humans and mice to measure reversible biological age changes in response to various stressful stimuli. We find that biological age may increase over relatively short time periods in response to stress, but this increase is transient and trends back toward baseline following recovery from stress. Using stressful events to investigate this question, we further find that second generation human DNAm clocks give consistent outputs, whereas first generation human DNAm clocks generally lack the sensitivity to detect transient changes in biological age. Finally, using COVID-19 as a model of severe infectious disease that triggers a reversible increase in biological age, we demonstrate that recovery of biological age following a stress-induced increase is a useful model with which to predict potential anti-aging drugs. Overall, our data suggest that increases in biological age due to stress may be an actionable target for future antiaging interventions.

## Results

### Heterochronic parabiosis induces a reversible increase in biological age

We began to examine possible fluctuations in biological age by using a mouse model of heterochronic parabiosis.^19,20^ We have previously evaluated the effect of this procedure on the old parabionts.^20^ Here, we tested whether exposure of young mice to aged circulation would induce a change in biological age, and whether such a change is reversible. We surgically joined pairs of either 3-month-old mice (isochronic) or a 3-month-old mouse and a 20-month-old mouse (heterochronic). After three months of parabiosis, the pairs were separated and allowed to recover for two months (Fig. 1a). Livers from the young mice were then analyzed using DNAm clocks, adjusting for chronological age (see Methods). The resulting *DNAm age acceleration* parameter (i.e., chronological-age-adjusted DNAm age) allows for unbiased statistical comparisons between age/treatment groups. This is particularly important for human datasets (below) where samples originate from subjects of diverse chronological age.

**Fig. 1.**
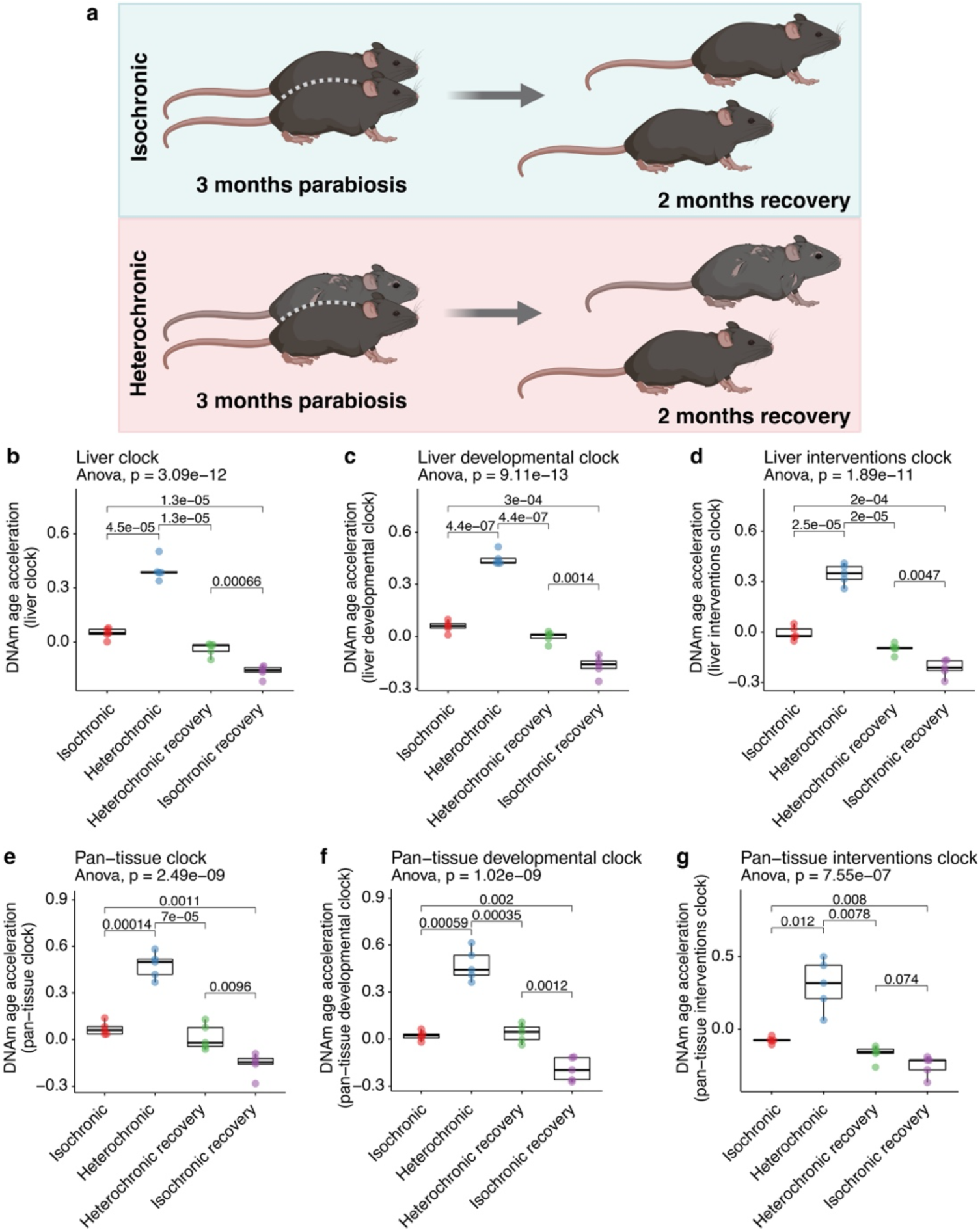
Young mice exposed to aged circulation undergo a reversible increase in biological age. (a) Setup of parabiosis experiment. Young (3-month-old) mice were surgically joined with either another young mouse (isochronic) or an old (20-month-old) mouse (heterochronic) for 3 months. Following the parabiosis period, mice were separated and allowed to recover for a further 2 months. Livers from young mice were analyzed using DNAm clocks to assess biological age. (b–g) DNAm age acceleration results from the HorvathMammalMethyl40 liver (b), liver developmental (c), liver interventions (d), pan-tissue (e), pan-tissue developmental (f), and pantissue interventions (g) clocks. P values were calculated with ANOVA and unpaired t-tests. Sample sizes: b–g, n= 6 for isochronic and isochronic recovery, and n=5 for heterochronic and heterochronic recovery.

We first analyzed methylation data profiled by the HorvathMammalMethylChip40, which reports the methylation status of approximately 36,000 CpG sites conserved across 159 mammalian species.^21^ Compared to isochronic controls, three liver-specific DNAm clocks revealed a significant increase in biological age of heterochronic parabionts of about 10-fold on average (Fig. 1b–d). Remarkably, biological age of heterochronic parabionts returned to baseline following detachment and recovery. Interestingly, we also observed a small but significant decrease in biological age of isochronic parabionts following recovery. We attribute this decrease to recovery from the stress of the surgery and parabiosis procedure.

Pan-tissue clocks (i.e., clocks trained on and applicable to multiple tissues^9^) fully recapitulated the effects we observed using liver-specific clocks (Fig. 1e–g). Furthermore, universal mammalian clocks trained on methylation data from 185 mammalian species^22^ also recapitulated our observations, although these changes did not rise to the level of statistical significance in all cases (Supplementary Fig. 1a–b). Nevertheless, these results indicate that biological age of young mice can be reversibly increased by heterochronic parabiosis.

To substantiate these findings, we subjected liver samples from our parabiosis animals to reduced representation bisulfate sequencing (RRBS) and applied several epigenetic clocks trained on RRBS data. We found good agreement with the methylation microarray clocks above. The Meer *et al*.^10^ and Petkovich *et al*.^11^ clocks, previously developed by our laboratory, fully recapitulated the reversible increase in biological age in heterochronic parabionts, as did the Stubbs *et al*.^12^ clock (Supplementary Fig. 1c–e). Biological ages of isochronic vs. heterochronic parabionts were not significantly different in the Wang *et al*. clock,^3^ although the reversal of biological age in heterochronic parabionts after recovery was significant (Supplementary Fig. 1f). The Thompson *et al*. clock^13^ indicated a significant difference in biological age between isochronic and heterochronic parabionts, but no significant DNAm age reversal following recovery (Supplementary Fig. 1g). Overall, the trends in the data were highly consistent in all clocks across both methylation profiling platforms. Thus, heterochronic parabiosis induces an increase in biological age of the young parabiont that is reversed following separation and recovery.

### Trauma surgery induces a reversible change in biological age of elderly patients

Having demonstrated that a transient increase in biological age can be experimentally induced, we sought to identify “natural” situations that similarly cause a reversible change in biological age. Given the links between chronic stress and accelerated biological age,^23,24^ we hypothesized that an acute, highly stressful health event might induce such a change. To test this hypothesis, we examined DNAm of blood samples from elderly patients undergoing major surgery.^**25**^ Blood from these patients was taken at three points: (1) immediately before surgery; (2) the morning after surgery; and (3) 4–7 days post-surgery, before discharge from the hospital. Using second-generation human DNAm clocks (DNAmPhenoAge,^14^ DNAmGrimAge,^15^ and DunedinPACE^17^), we found a significant increase in biological age of patients undergoing emergency surgical repair of a traumatic hip fracture. Remarkably, this increase occurred in under 24 hours, and biological age returned to baseline 4–7 days post-surgery (Fig. 2a–c). Interestingly, two other non-trauma surgeries did not produce this effect. Patients undergoing elective hip surgery showed an overall increase in biological age following surgery, approaching an age acceleration of 0 (or a DunedinPoAm score of just over 1) by the end of their hospitalization (Fig. 2d–f). We note that these patients started at a lower biological age relative to emergent patients (around -5 age acceleration for DNAmPhenoAge and DNAmGrimAge, and 1 for DunedinPoAm), likely reflecting selection of otherwise healthy surgical candidates and preoperative preparation for a planned surgery.^26^ Finally, patients undergoing elective colorectal surgery showed no significant changes in biological age over the course of their care, likely reflecting the less intensive nature of such surgeries (Fig. 2g–i).

**Fig. 2.**
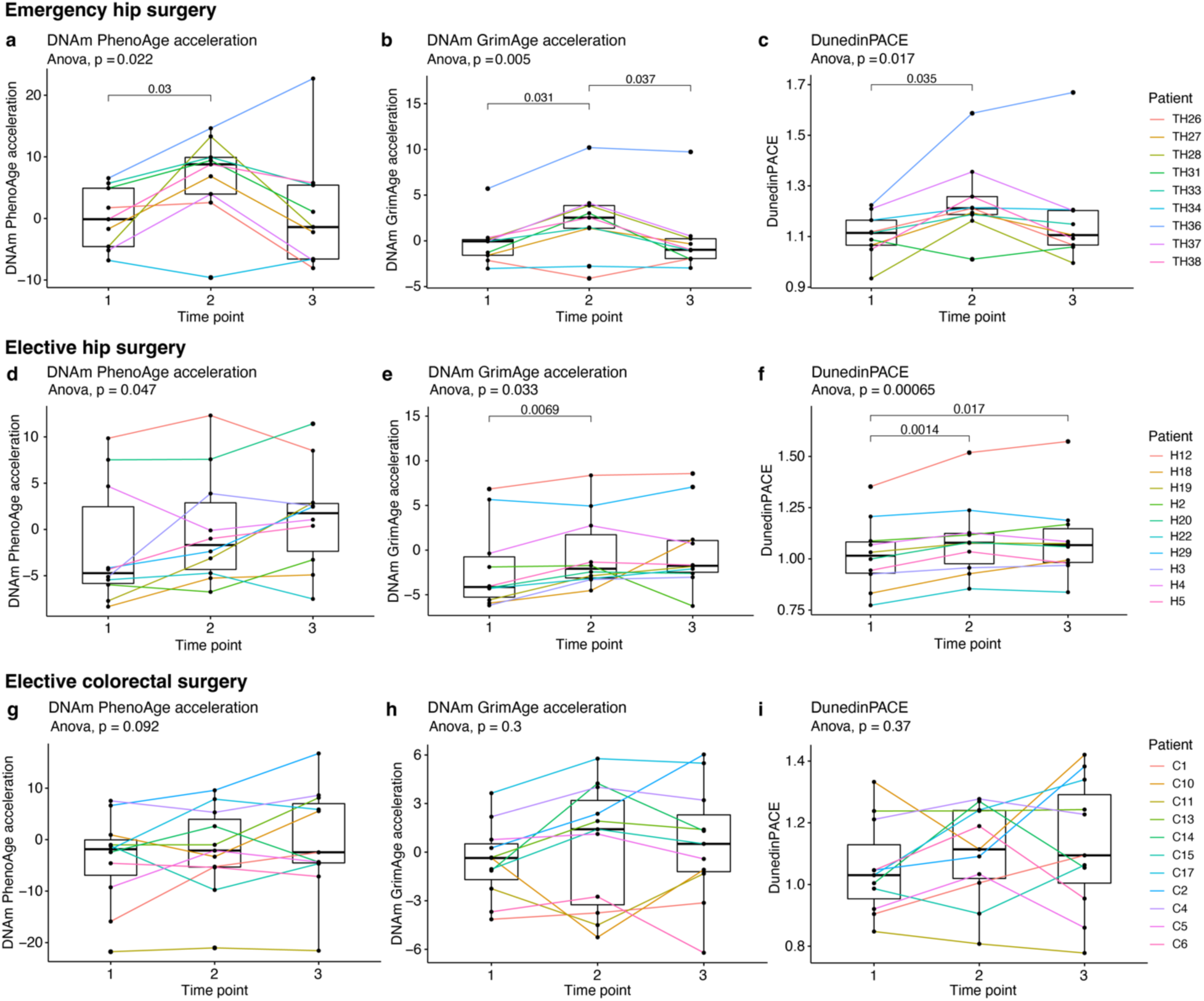
Patients undergoing major emergency (but not elective) surgery experience a reversible increase in biological age. (a–c) Second-generation DNAm age biomarkers for patients undergoing emergency surgery to repair traumatic hip fractures determined using DNAmPhenoAge (a), DNAmGrimAge (b), and DunedinPoAm38 (c). (d–f) As above, but for patients undergoing elective hip surgery. (g–i) As above, but for patients undergoing elective colorectal surgery. In all panels, time point 1 corresponds to immediately before surgery; time point 2 corresponds to the morning after surgery; and time point 3 corresponds to the day of discharge from the hospital, 4–7 days post-surgery. P values were calculated with repeated-measures ANOVA and paired t-tests. Sample sizes: a–c, n=9; d–f, n=10; g–i, n=11.

In all three cohorts of surgical patients, first-generation DNAm clocks (Horvath DNAm age,^6^ Hannum DNAm age,^7^ and Skin & Blood DNAm age^8^) showed no significant changes (Supplementary Fig. 2). A recent preprint introduced principal component (PC) corrected versions of the major DNAm aging clocks to correct for technical noise and improve performance of the clocks on longitudinal data.^27^ Application of these PC clocks to our data yielded consistent results with the original versions of these clocks (Supplementary Fig. 3). In some cases, first-generation PC clocks revealed significant changes that agreed with second-generation clocks; however, second-generation PC clocks still showed more consistent significant changes overall (we explore this further in other datasets below). Most importantly, the trends we observed using the original clocks agreed with those revealed by PC clocks: emergency hip surgery patients underwent a reversible increase in biological age, elective hip surgery patients started at negative age accelerations and underwent a gradual increase towards baseline, and elective colorectal surgery had no effect on biological age.

We also utilized DNAm predictors of blood cell composition to analyze blood cell dynamics in this cohort of patients.^2,28^ For emergency hip surgery patients, we found significant differences in the counts of B cells, several subsets of T cells, plasmablasts, natural killer (NK) cells, and monocytes. Elective hip surgery patients experienced significant fluctuation in levels of plasmablasts, NKs, and Monocytes, and elective colorectal surgery patients showed no significant fluctuations in any of the cell types analyzed (Supplementary Fig. 4).

### Biological age of mice reversibly increases during pregnancy

We further examined reversible changes in biological age by testing the effect of pregnancy, given the significant biological overlap between pregnancy and aging. Pregnancy is a highly physiologically stressful event, with nearly every organ system subject to increased demand to support the developing fetus.^29^ Because of the resulting damage accumulation and increased incidence of age-related diseases such as diabetes and heart disease, pregnancy has even been suggested as a model for aging.^30^ With these considerations in mind, we hypothesized that biological age would increase over the course of pregnancy and return to baseline following delivery.

We began by studying a mouse model of pregnancy. Following a baseline blood sample, C57Bl/6 mice were mated, and two blood samples were taken during the early and late phases of their pregnancies. Following parturition and a period of recovery, a final blood sample was collected (Fig. 3a). We subjected DNA isolated from these blood samples to the HorvathMammalMethylChip40 for methylation profiling. The mouse blood clock revealed a significant decrease in biological age following parturition (Fig. 3b), but no change in age-matched mice that were also mated but did not become pregnant (Fig. 3c). Interestingly, the blood developmental clock showed an increase in biological age after mice became pregnant that resolved following parturition and recovery (Fig. 3d). Again, no change was detected by this clock in non-pregnant animals (Fig. 3e). We suspect that since the developmental clock was built using CpGs whose methylation levels change during development, this clock may be more suitable to evaluate pregnancy, a developmentally relevant process. In any event, taking the two clocks together, we conclude that pregnancy may induce a reversible increase in biological age.

**Fig. 3.**
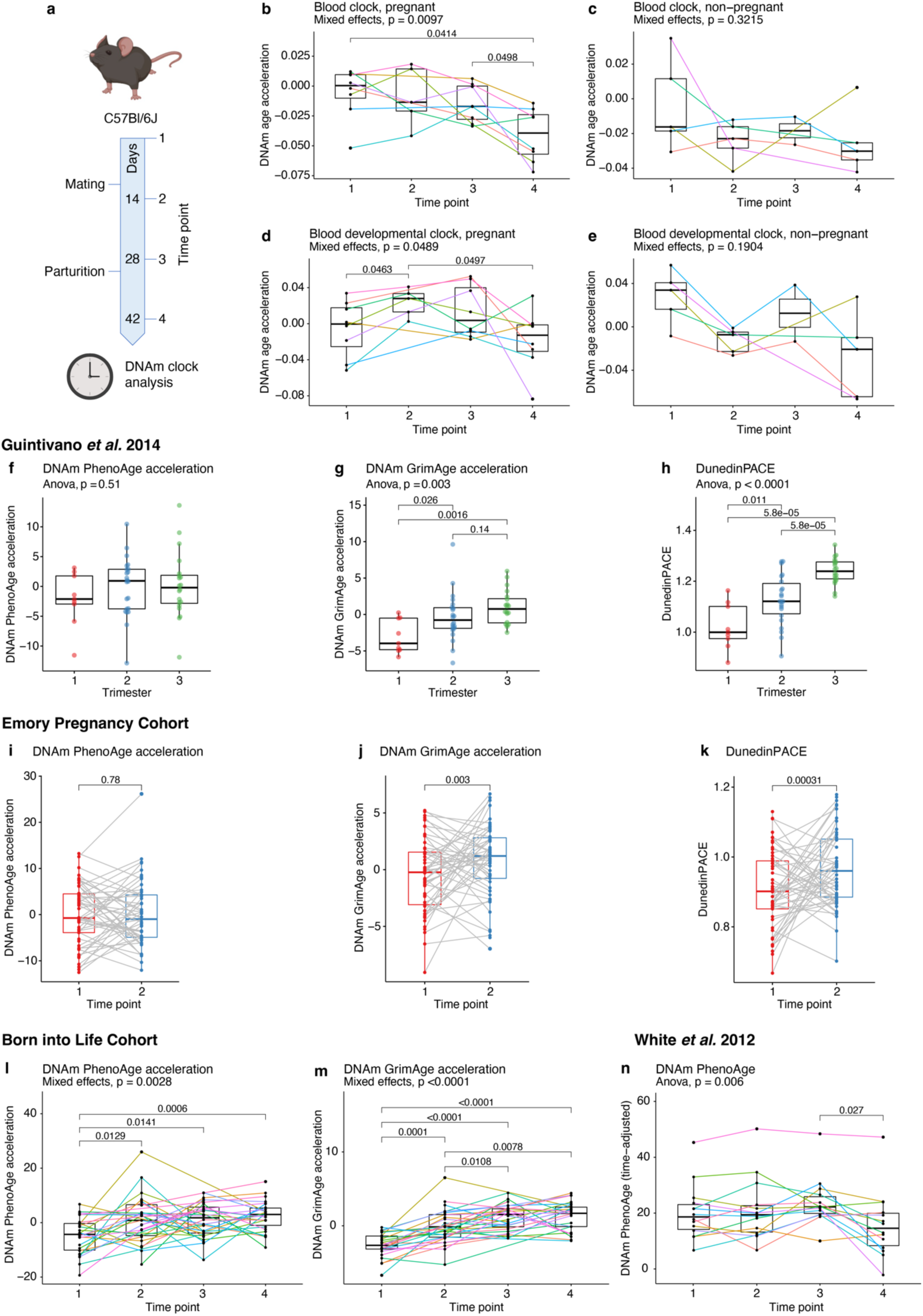
Mice and humans experience an increase in biological age over the course of pregnancy that is reversed following parturition. (a) Timeline of mouse pregnancy study. Note that “days” here refers to experimental days, not embryonic ages. Blood was collected from C57Bl/6 mice before, during, and after pregnancy, and DNA isolated from this blood was subjected to DNAm clock analysis. (b–c) Blood clock DNAm age acceleration results from pregnant (b) and non-pregnant (c) mice. (d–e) Blood developmental clock DNAm age acceleration results from pregnant (d) and non-pregnant (e) mice. (f–h) Cross-sectional DNAm age acceleration analysis of pregnant Americans across the three trimesters of pregnancy using DNAmPhenoAge (f), DNAmGrimAge (g), and DunedinPoAm38 (h). (i–k) DNAm age biomarkers (as in f–h) for a longitudinal study of pregnant African Americans with two blood samples collected over the course of pregnancy. Time point 1 corresponds to 7–15 weeks of pregnancy; time point 2 corresponds to 24–32 weeks of pregnancy. (l–m) DNAmPhenoAge (l) and DNAmGrimAge (m) acceleration results from Swedish mothers longitudinally tracked over the course of pregnancy. Time point 1 corresponds to pre-pregnancy; time point 2 corresponds to 10–14 weeks of pregnancy; time point 3 corresponds to 26–28 weeks of pregnancy; time point 4 corresponds to 2– 4 days postpartum. (n) DNAmPhenoAge (adjusted for the passage of time; see Methods for details) for a cohort of American mothers longitudinally tracked over the course of pregnancy and postpartum. Time point 1 corresponds to early pregnancy; time point 2 corresponds to mid-pregnancy; time point 3 corresponds to delivery; time point 4 corresponds to 6 weeks postpartum. P values were calculated using either repeated-measures ANOVA and paired t-tests or a mixed effects model with post-hoc pairwise comparison testing (see Methods). Sample sizes: b and d, n=8; c and e, n=5; f–h, n=9, 22, and 20 for trimesters 1, 2, and 3, respectively; i–k, n=53; l–m, n=33 total subjects who each provided up to 4 samples; n, n=14. Note that for the Born into Life Cohort, due to data sharing limitations, we were unable to obtain the CpG data necessary to analyze DunedinPACE. Note also that the White *et al*. 2012 dataset was generated using the Illumina HumanMethylation27 Beadchip, which limited our analysis to DNAm PhenoAge (panel n) and Horvath DNAm age (Supplementary Fig. 5j).

### Human pregnancy causes a reversible increase in biological age

To corroborate and expand on our results in mice, we analyzed methylation datasets from several cohorts of pregnant humans (Supplementary Table 1). Most available longitudinal methylation datasets tracking women over the course of pregnancy cover the period from pre-/early pregnancy up to (or very shortly after) delivery. A cross-sectional dataset^31^ of 54 pregnant American women from whom blood was sampled at during one trimester of their pregnancy showed no difference in DNAmPhenoAge acceleration (Fig. 3f), but a significant increase in biological age from the first to third trimesters was found using DNAmGrimAge (Fig. 3g), and DunedinPACE revealed significant increases between both the first to second and second to third trimesters (Fig. 3h). Similarly, DNAmGrimAge and DunedinPACE, but not DNAmPhenoAge, revealed an increase in biological age from early to late pregnancy in a longitudinal dataset^32^ consisting of African American women who each provided two blood samples over the course of their pregnancies (Fig. 3i–k). In a cohort of pregnant Swedish women,^33,34^ both DNAmPhenoAge and DNAmGrimAge revealed a progressive increase in biological age from pre-pregnancy (time point 1) to 2–4 days postpartum (time point 4) (Fig. 3l–m). Thus, biological age increases in human pregnancy up to the point of parturition, consistent with the effects we found in mice. As with our human surgery data analysis (Supplementary Fig. 2), first generation clocks did not detect any changes in these pregnancy datasets (Supplementary Fig. 5a–i). PC versions of first-generation clocks also did not detect significant changes in any of the datasets to which we were able to apply them (Supplementary Fig. 6). PC versions of second-generation clocks yielded consistent results with original clocks (Supplementary Fig. 6).

We identified one dataset^**35**^ in which women were tracked longitudinally over the course of their pregnancy through 6-weeks postpartum. However, methylation profiling for this cohort was performed using the Illumina HumanMethylation27 beadchip, which limited the number of clocks we could apply to the Horvath multi-tissue clock and DNAmPhenoAge. DNAmPhenoAge (corrected for the passage of time, as chronological ages were not available for this dataset; see Methods) revealed a trend toward higher biological age at delivery, followed by a significant reversal of biological age at 6 weeks postpartum (Fig. 3n), consistent with our mouse data above. Horvath DNAm age closely mirrored the overall trend but did not rise to the level of statistical significance (Supplementary Fig. 5j). Analyses of blood cell composition predicted from methylation data for the human pregnancy cohorts did not reveal consistent changes in cell composition over the course of pregnancy between datasets (Supplementary Fig. 7). Thus, it is unlikely that changes in blood composition alone can explain the highly consistent effects we observe on biological age. Taking all our analyses in mice and humans together, we conclude that pregnancy induces a reversible increase in biological age, peaking around delivery and resolving postpartum.

### Severe COVID-19 causes a reversible increase in biological age

We next hypothesized that severe infectious disease might cause reversible changes in biological age. COVID-19 is an ideal test case given its strong links to aging.^36^ Not only are the elderly up to 90-fold more vulnerable to death from COVID-19,^37^ but we and others have previously reported that accelerated biological age is associated with incidence and severity of COVID-19.^38–42^ Longitudinal biological age data covering the COVID-19 disease course is also extremely limited. A recent report included a subset of longitudinal samples, but the sample size (n=3) precluded any statistical analysis.^42^ We therefore sought to investigate whether COVID-19 induces a reversible change in biological age.

To directly test how biological age changes over the course of severe infectious disease, we obtained longitudinal blood samples from COVID-19 patients. Our cohort consisted of patients who tested positive for COVID-19 by RT-PCR, were admitted to an intensive care unit, survived the disease, and provided multiple blood samples spanning the course of their hospitalization. Because the patients in our cohort were generally already admitted to the ICU by the time the first available blood sample was taken, we hypothesized that the major effect we would observe was a reversal of *already* accelerated biological age. Given the known differences in both disease course and outcomes in males and females (with males generally experiencing poorer outcomes^43^), we separated our analysis by sex.

DNAmPhenoAge indicated a significant reversal of biological age in females following discharge from the ICU (i.e., time points 3–4), but no significant change in males (Fig. 4a). Similarly, DNAmGrimAge indicated an increase in biological age that was partially reversed by the time of ICU discharge for females. This was marginally significant overall, and no significant change was observed in males (Fig. 4b). In both cases, we note that male patients exhibited much more heterogeneity in the trajectories of their biological age over the disease course.

**Fig. 4.**
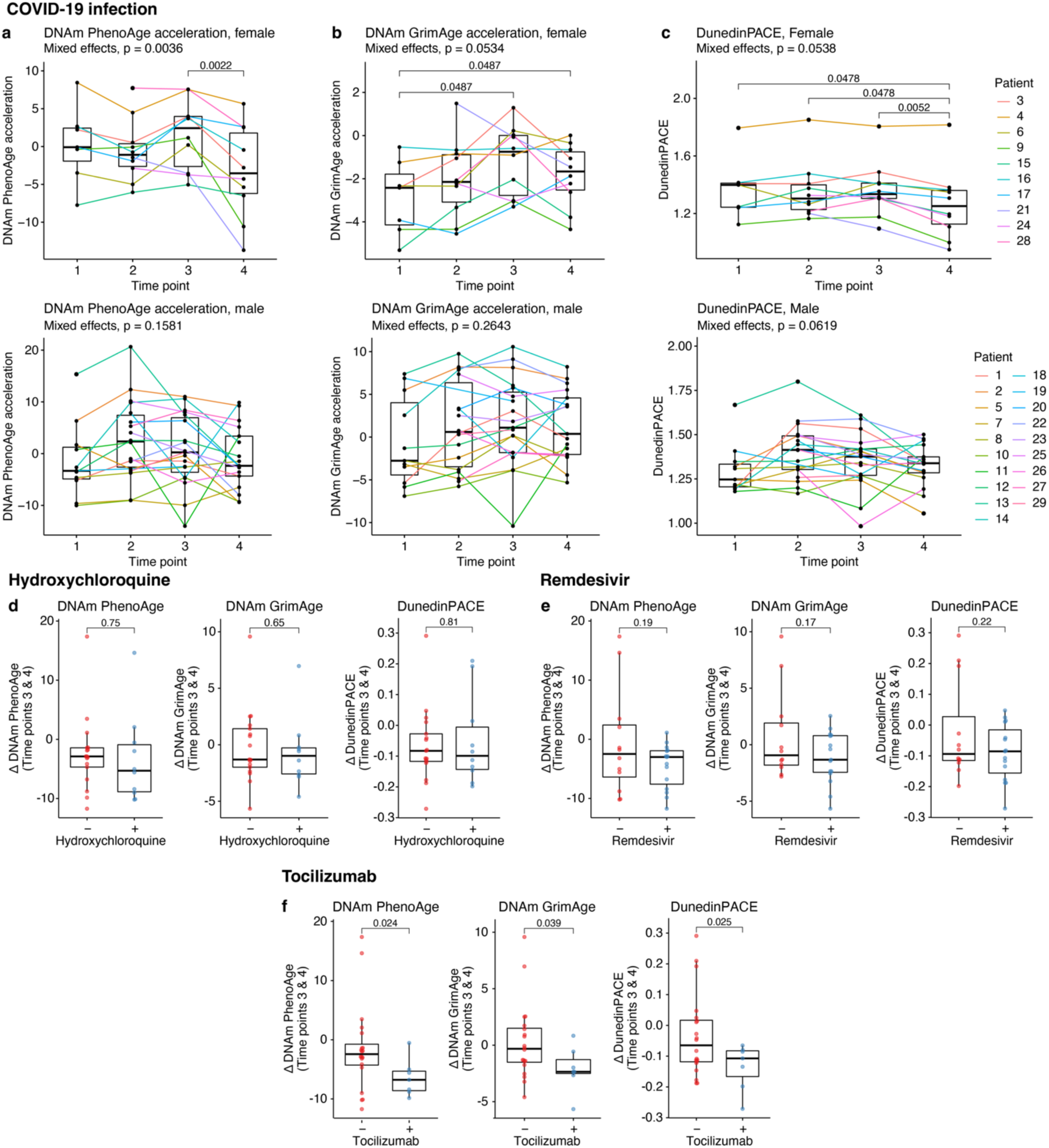
Patients with severe COVID-19 experience a reversible increase in DNAm age; treatment with tocilizumab enhances DNAm age recovery following ICU discharge. (a–c) DNAm age acceleration results for DNAmPhenoAge (a), DNAmGrimAge (b), and DunedinPoAm38 (c). All upper panels show data for female patients and all lower panels show data for male patients. Time point 1 is within 5 days of ICU admission; time point 2 is within 5 days of the midpoint of the ICU stay; time point 3 is within 5 days of the date of discharge from the ICU; timepoint 4 is ≥7 days post-ICU discharge. (d–f) DNAm age recovery, defined as the difference in DNAm age acceleration between time points 3 and 4), for patients treated with hydroxychloroquine (d), remdesivir (e), or tocilizumab (f). In a–c, p values were calculated using a mixed effects model with post-hoc pairwise comparison testing. In d–f, p values were calculated with unpaired t-tests. Sample sizes: a–c, n=10 female and n=19 male subjects total who each provided up to 4 samples; d, n=19 untreated and 10 treated patients; e, n=12 untreated and 17 treated patients; f, n=21 untreated and 8 treated patients.

In the case of DunedinPACE (in which the normal pace of aging is 1), we found that the pace of aging was already elevated by ~25% by the point of ICU admission (time point 1) for both sexes. This was reversed following discharge from the ICU, although not fully to baseline (Fig. 4c). As in all cases involving human samples above, first generation clocks did not detect any changes in either males or females (Supplementary Fig. 8). PC clock results generally did not rise to the level of statistical significance, although trends for second-generation PC clocks were consistent with the original second-generation clocks (Supplementary Fig. 9). Few types of blood cells showed significant variation over the course of the disease, and those that did (CD4^+^ T cells, plasmablasts, NKs, granulocytes) did not change consistently between male and female patients (Supplementary Fig. 10). On the whole, we conclude that a severe infectious disease such as COVID-19 can induce a reversible increase in biological age, although the results are nuanced and seem to be both sex and clock-specific.

### Reversal of elevated biological age can be used to predict anti-aging interventions

The observed reversal in biological age of COVID-19 patients following discharge from the ICU provides a tool with which to predict interventions that may potentially allow patients to recover their biological age more rapidly following a stressful event. We thus investigated the effect of experimental interventions received by our COVID-19 patient cohort on their ability to reverse their increased biological age. Largely due to the timeframe during which these samples were collected (March–June 2020), this group of interventions included hydroxychloroquine, remdesivir, and tocilizumab.^44^ We calculated biological age recovery by subtracting biological age at time point 4 (≥7 days after ICU discharge) from time point 3 (ICU discharge). Neither the antimalarial hydroxychloroquine nor the broad-spectrum antiviral remdesivir showed any effects on biological age recovery (Fig. 4d–e). Interestingly, however, patients treated with tocilizumab, a monoclonal antibody targeting the interleukin-6 receptor, showed a greater recovery of biological age (by all three second-generation clocks) than patients that did not receive this intervention (Fig. 4f). Thus, tocilizumab may warrant further investigation as a geroprotective drug.

## Discussion

This study reveals that biological age of humans and mice is not static nor steadily increasing but undergoes reversible changes over relatively short time periods of days–months according to multiple independent epigenetic aging clocks. This finding of fluid, fluctuating, malleable age challenges the longstanding conception of a unidirectional upward trajectory of biological age over the life course. Previous reports have hinted at the possibility of short-term fluctuations in biological age,^45^ but the question of whether such changes are reversible has, until now, remained unexplored. Critically, the triggers of such changes were also unknown. We established that a reversible biological age change can be experimentally induced in animals subjected to heterochronic parabiosis. An increase in biological age upon exposure to old blood is consistent with previous reports of detrimental age-related changes upon heterochronic blood exchange procedures.^46–48^ However, reversibility of such changes, as we observed (Fig. 1 and Supplementary Fig. 1), has not yet been reported. From this initial insight, we hypothesized that other naturally occurring situations might also trigger reversible changes in biological age.

A clear pattern that emerged over the course of our studies is that exposure to stress increased biological age. When the stress was relieved, biological age could be fully or partially restored. This is perhaps most clearly demonstrated by our analysis of biological age changes in response to major surgery (Fig. 2). Although we did not observe the effect in the case of elective surgeries, where patients are pre-screened for surgical candidacy and advised to follow strict preparation guidelines^26^ (likely reflected in the lower biological age found in these patients on the day of their surgery), we saw a strong and rapid increase in biological age in trauma patients following emergency surgery. This increase was reversed, and biological age was restored to baseline in the days following the surgery. Of note, the patients in this cohort were elderly (mean age 77.9 years) implying, surprisingly, that even people of advanced chronological age have the capacity to reverse a stress-induced increase in their biological age. Reversible changes in biological age were also found in response to pregnancy and COVID-19, implying that such changes may be rather common responses to stress. These situations (and others yet to be discovered) that trigger a rapid increase in biological age are likely good candidate models for testing the ability of anti-aging drugs to improve clinical outcomes. Moreover, our finding that biological age reversal is achievable on the scale of days (Fig. 2a–c, cf. time points 2–3) strongly points to the potential utility of anti-aging drugs in diseases/medical interventions that lead to increased stress, such as major surgery. The ability of tocilizumab to enhance the biological age recovery of convalescent COVID-19 patients (Fig. 4f) lends further credence to this notion.

From a technical standpoint, across human data sets examined, we consistently observed that first generation DNAm clocks were not able to detect significant effects found by second generation clocks applied to the same data, even after PC correction in nearly all cases.^27^ This may imply that second generation clocks intrinsically possess features that render them more sensitive to short term fluctuations in biological age. For instance, the integration of multiple age-related biomarkers into the models of second-generation clocks may render them more sensitive to transient fluctuations in biological age compared to first generation clocks, which are trained only on chronological age. Whatever the underlying reason, this highlights the critical importance of judicious selection of DNAm clocks appropriate to the analysis at hand, especially in light of the many clocks continuously coming to the fore. Nevertheless, we obtained consistent outputs across second-generation clocks applied across our human DNAm datasets, as well as agreement with mouse models in the case of pregnancy, bolstering our confidence in our conclusions.

While this study highlights a previously unappreciated aspect of dynamic biological aging, we acknowledge some important limitations. First, we focused our assessment of biological age to DNAm clocks, the most powerful aging biomarker currently available. It is our hope that as the ongoing expansion of the aging biomarkers field proceeds, additional biomarkers that rival or exceed the power of DNAm clocks will allow us to confirm our conclusions using orthogonal approaches to measure biological age. Indeed, recent analyses of complete blood counts and physical activity are consistent with our findings of fluctuations in biological age across the entire life.^45^ Second, a concern common to all studies that utilize biomarkers of aging is discrimination of true effects from artifacts of the biomarkers. Although no biomarker is perfect, several lines of evidence give us confidence that our observations represent true modulations of biological age: (1) where we were able to analyze across species, the effects were consistent; (2) we observe effects consistently with one class of DNAm biomarkers (second-generation clocks), but not another (first-generation clocks). We would expect artifactual “positive” results to occur randomly across the biomarkers analyzed; and (3) several distinct models—surgery, pregnancy, and COVID-19—united by the severe physiological stress they induce, caused similar effects on the biomarkers. Future work will be needed to link, for instance, successful recovery of biological age following a stressful event to improved clinical outcome.

In the most fundamental sense, our data reveal the dynamic nature of biological age: stress can trigger a rapid increase in biological age, and such a change can be reversed. Importantly, this implies both the existence of intrinsic mechanisms to reverse increased biological age and the opportunity to reverse transient increases in biological age therapeutically. The findings also imply that severe stress increases mortality, at least in part, by increasing biological age. This notion immediately suggests that mortality may be decreased by reducing biological age, and that ability to recover from stress may be an important determinant of successful aging and longevity. Finally, biological age may be a useful parameter in assessing physiological stress and its relief.

## Methods

### Mouse experiments

All mouse experiments were approved by the Mass General Brigham IACUC or the Duke University IACUC. C57Bl/6 mice were obtained from Jackson Laboratories and acclimated to our animal facility for at least 48 h before being subjected to any experimental manipulation. Aged C57Bl/6 mice for parabiosis experiments were obtained from the NIA aged rodent colony. Mice were maintained in a barrier facility in sterilized, ventilated cages and fed standard laboratory chow (LabDiet 5053) and reverse osmosis drinking water ad libitum and maintained on a 12h:12h light:dark cycle. Mice were generally housed socially (5 mice/cage) except for the pregnancy studies wherein male mice were housed individually after mating. Mice were humanely euthanized at the conclusion of each experiment by CO2 exposure followed by cervical dislocation.

### Mouse parabiosis experiments

Parabiosis was carried out as previously described.^20^ Female C57Bl/6 mice were pre-screened to minimize body size differences, and were randomly assigned to parabiosis pairs. Isochronic pairs consisted of two 3-month-old mice and heterochronic pairs consisted of one 3-month-old mouse and one 20-month-old mouse. Pairs were surgically attached and maintained for 3 months. Following 3 months of parabiosis, a subset of mice were euthanized for analysis and another subset were surgically separated. Fascia and skin were sutured closed following separation, and mice were allowed to recover for 2 months, after which they were for euthanized for analysis.

### Mouse pregnancy experiments

C57Bl/6 mice (11 weeks old) were obtained from Jackson Laboratories. Three days before mating, male mice were separated into individual cages and soiled bedding from male cages was added to female cages to induce estrus.^50^ 1:1 mating pairs were set up in the evening and left overnight. Females were removed from male cages the following morning and inspected for copulatory plugs. Pregnant females were identified by daily tracking of body weight. Blood was collected in EDTA tubes by submandibular vein puncture every two weeks to create a series of four samples per mouse: (1) 10 days before mating; (2) 4 days after mating; (3) 18 days after mating; and (4) 32 days after mating, generally corresponding to ~2 weeks postpartum. Pups were humanely euthanized shortly after birth allowing mothers to recover from pregnancy without needing to nurse. Blood was snap-frozen in liquid nitrogen immediately after collection and stored at –80°C until needed.

### COVID-19 study

This study was approved by the Mass General Brigham IRB (protocol number 2020P004121). We selected a cohort of patients with RT-PCR-confirmed COVID-19 who were admitted to the intensive care units of Brigham and Women’s Hospital (Boston, MA, USA). Clinical blood samples from these subjects were obtained through the Crimson Core facility of Mass General Brigham. Buffy coats from these blood samples were used as a source of DNA for methylation profiling as described elsewhere. Clinical and demographic data were collected by review of electronic medical records.

### Isolation of genomic DNA

DNA was isolated from human buffy coat samples at the Crimson Core facility (Mass General Brigham) using the QIAamp DNA Blood Mini Kit (Qiagen 69504) following the manufacturer’s protocol. Eluted DNA was concentrated using a speedvac. DNA was isolated from mouse tissues using the DNeasy Blood and Tissue Kit (Qiagen) following the manufacturer’s protocol. Generally, 50–100 μl of blood or ~25 mg of solid tissue was used as starting material. Concentration of DNA samples was determined using the Qubit dsDNA BR assay kit (Invitrogen). Isolated DNA was stored at –20°C.

### DNA methylation profiling

Methylation data was generated through the Epigenetic Clock Development Foundation. Human DNA samples for the COVID study were subjected to the Infinium MethylationEPIC array (Illumina) at AKESOgen Inc., and mouse DNA samples were subjected to the HorvathMammalMethylChip40 at the UCLA Neuroscience Genomic Core (UNGC). Samples were randomized to avoid introduction of batch/chip effects, but longitudinal samples from a single patient/mouse were run on the same chip. All sample preparation/processing was carried out according to the Illumina kit protocols.

### Other sources of human methylation data

Illumina HumanMethylation450 BeadChip data for surgical patients from Sadahiro *et al*.^25^ were downloaded from GEO (GSE142536). Illumina HumanMethylation450 BeadChip data for the Emory University African American Microbiome in Pregnancy Cohort are publicly available via GEO (GSE107459). In our study, only paired samples were analyzed; participants with only a single blood sample were excluded. Illumina HumanMethylation450 BeadChip data from Guintivano *et al*.^31^ are publicly available via GEO (GSE44132). Chronological age data for this cohort were kindly provided by Prof. Zachary Kaminsky (The Royal, Canada). Illumina MethylationEPIC BeadChip data from the Born into Life cohort^33,34^ were kindly provided by Prof. Catarina Almqvist (Karolinska Institutet, Sweden). Detailed information for these datasets can be found in Supplementary Table 1.

### DNAm clock analysis

For mammalian microarray analysis, raw methylation data were first normalized using the SeSAMe R package and beta values were calculated. DNAm age biomarkers were calculated as previously described.^9^ For publicly available human datasets, if raw idat methylation files were available, they were processed using the minfi R package.^51^ Data were first preprocessed using noob normalization and then beta values were calculated using the getBeta function. For datasets where only raw or normalized methylation data/calculated beta values were the only data available, these data were used directly. Human DNAm age biomarkers were calculated using the online Hovath DNA Methylation Age Calculator,^52^ which calculates Horvath DNAm age,^6^ Hannum DNAm age,^7^ Skin & Blood DNAm age,^8^ DNAmPhenoAge,^14^ and DNAmGrimAge,^15^ among other parameters. DunedinPACE^17^ was calculated with the DunedinPACE R package.^53^ RRBS-based epigenetic clocks were applied as previously described.^10,11,20^ PC clocks were applied using publicly available code.^27,54^

### Data analysis

All DNAm age biomarkers were adjusted by chronological age to yield an age acceleration parameter. For human studies, this was carried out by calculating residuals from regressing DNAm age on chronological age. Note that this correction is neither necessary nor relevant for DunedinPACE. For mouse experiments, linear regressions are skewed by the strong effects of the experimental interventions applied (e.g. parabiosis). Thus, for mouse experiments, we calculated age acceleration by subtracting chronological age from DNAm age. In one case (the White *et al*. human pregnancy dataset), chronological ages were not available. To account for the passage of time in this longitudinal dataset, we corrected the DNAm age predictions by the average amount of time between each sample collection point, based on the methods of the original publication^35^ and the assumption that most women do not learn they are pregnant for 6 weeks on average. This corresponded to the following time corrections: 0.26, 0.635, and 0.75 years, respectively, for time points 2, 3, and 4.

### Statistics

For longitudinal datasets, repeated measures ANOVA or mixed effects models were first used to test for significant variance between time points. If these tests revealed a significant effect, paired t tests corrected by controlling the false discovery rate using the Benjamini-Hochberg method were carried out between groups. Exact p values are shown within all figures. For non-longitudinal datasets, a similar procedure was used except that traditional ANOVAs and unpaired t tests were used. All t tests were two-tailed. Sample sizes are indicated in figure legends.

## Acknowledgments

We thank Dr. Anastasia Shindyapina (Gladyshev Lab) and Nate Rogers (Brigham and Women’s Hospital Center for Comparative Medicine) for training in mouse handling and blood collection. We thank Lindsay Rutte, Tim Janicki, and Dr. Lynn Bry of the Brigham and Women’s Hospital Crimson Core Facility for assistance with COVID-19 sample acquisition. We thank Drs. Albert Higgins-Chen and Morgan E. Levine for sharing the PC clocks code prior to its publication. This study was funded by NIA grants (to VNG). JRP is supported by the BWH Organ Design and Engineering Training Program, NIBIB grant 5T32EB016652-07.

## Author contributions

Conceptualization: JRP, VNG

Methodology: JRP, BZ, CK, SHY, ATL, AH, JPW, SH, VNG

Investigation: JRP, BZ, CK, ATL, AH, SH

Essential data contribution: TG, AMH, EA, GP, CA

Visualization: JRP

Funding acquisition: JPW, VNG

Project administration: JPW, SH, VNG

Supervision: JPW, SH, VNG

Writing – original draft: JRP, VNG

Writing – review & editing: JRP, BZ, CK, SHY, ATL, AH, TG, AMH, EA, GP, CA, JPW, SH, VNG

## Data and materials availability

All data originally generated in this study will be made available in public databases upon publication. Unique materials will be made available upon reasonable request to the corresponding author.

