## Supplementary Information for "Biological age is increased by stress and restored upon recovery"

### Supplementary Figures

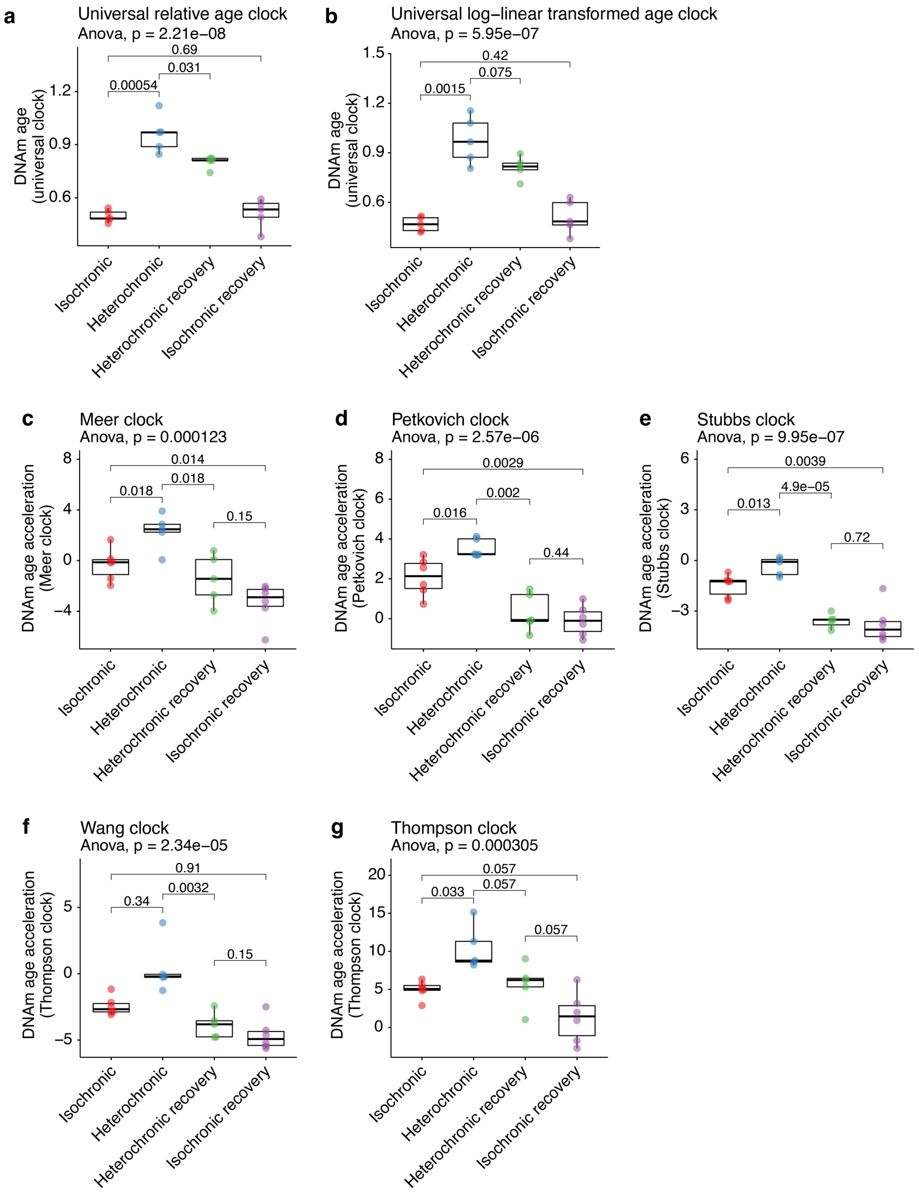

**Supplementary Figure 1. Universal mammalian and RRBS-based DNAm age biomarkers for mice subjected to parabiosis.** (a–b) DNAm age predictions from the universal pan-mammalian clock[1] for mice subjected to parabiosis, with age predictions based on relative age (a) or log-linear transformed age (b). (c–g) DNAm age acceleration results from RRBS-based DNAm clocks, including those of Meer *et al.*[2] (c), Petkovich *et al.*[3] (d), Stubbs *et al.*[4] (e), Wang *et al.*[5] (f), and Thompson *et al.*[6] (g). P values were calculated with ANOVA and unpaired t-tests. Sample sizes: a–g, n= 6 for isochronic and isochronic recovery, and n=5 for heterochronic and heterochronic recovery.

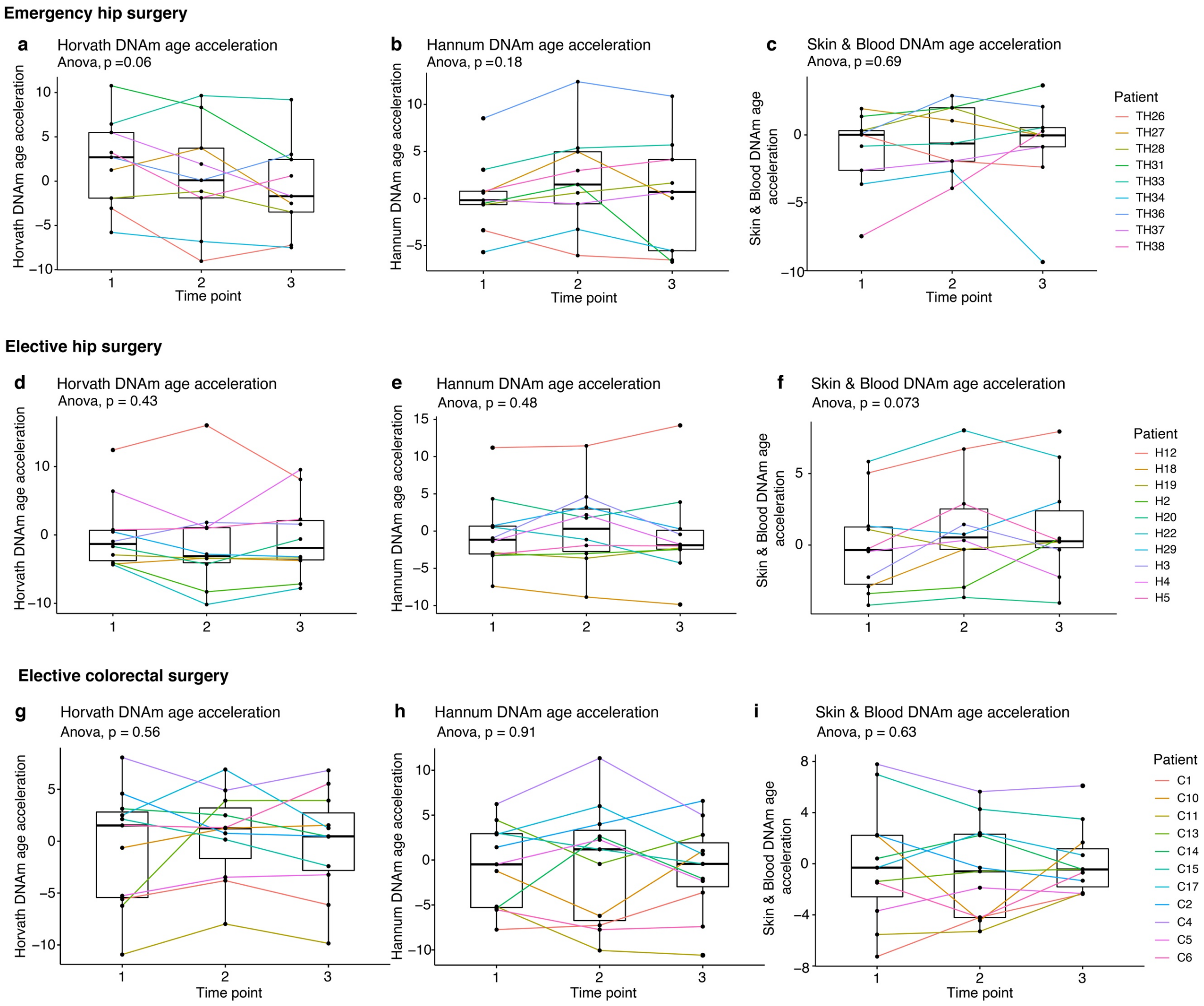

**Supplementary Figure 2. Horvath, Hannum, and Skin & Blood DNAm age acceleration for patients undergoing major surgery.** (a–c) First generation DNAm age acceleration results for patients undergoing emergency surgery to repair traumatic hip fractures calculated using Horvath DNAm age[7] (a), Hannum DNAm age[8] (b), and Skin & Blood DNAm age[9] (c). (d–f) As above, but for patients undergoing elective hip surgery. (g–i) As above, but for patients undergoing elective colorectal surgery. In all panels, time point 1 corresponds to immediately before surgery; time point 2 corresponds to the morning after surgery; and time point 3 corresponds to the day of discharge from the hospital, 4–7 days post-surgery. P values were calculated with repeated-measures ANOVA and paired t-tests. Sample sizes: a–c, n=9; d–f, n=10; g–i, n=11.

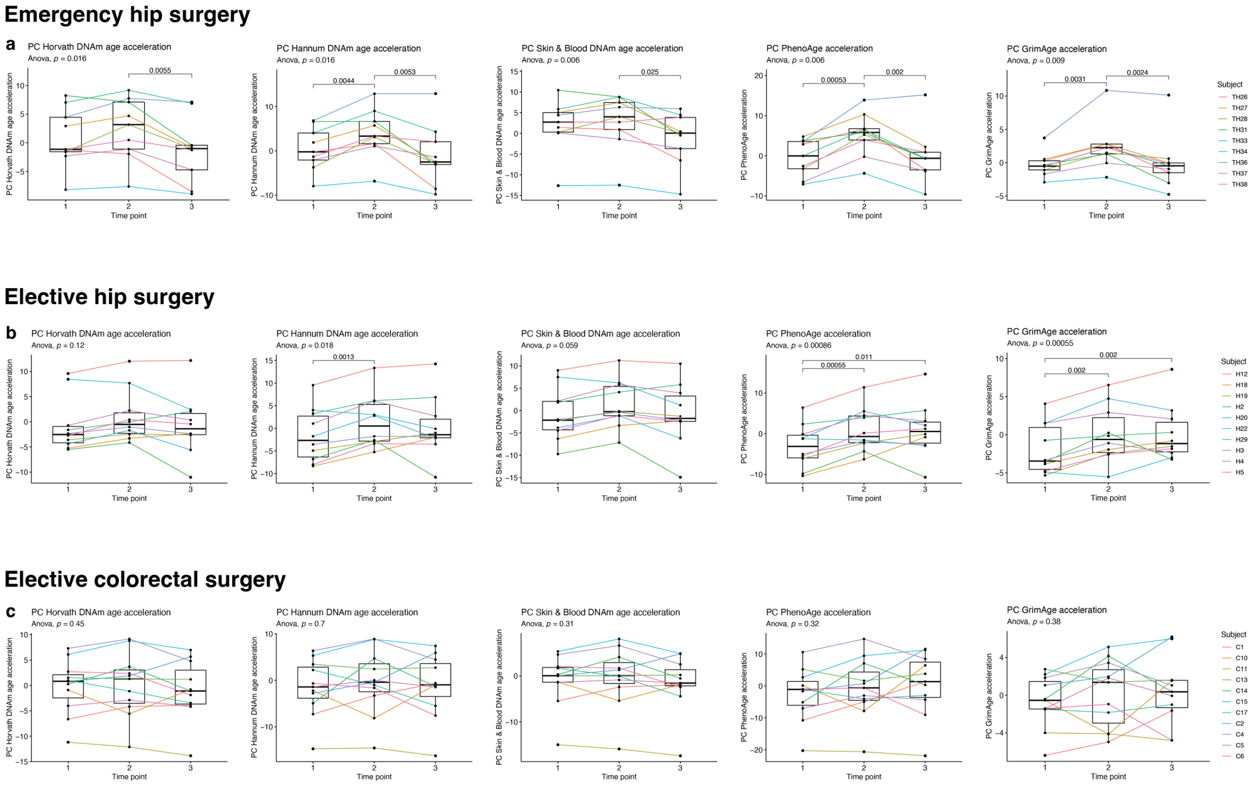

**Supplementary Figure 3. Principal component (PC) clocks results for patients undergoing major surgery.** (a–c) PC clocks (Horvath DNAm age, Hannum DNAm age, Skin & Blood DNAm age, DNAm PhenoAge, and DNAm GrimAge) [10] were applied to methylation data from patients undergoing emergency hip surgery (a), elective hip surgery (b), or elective colorectal surgery (c). P values were calculated with repeated-measures ANOVA and paired t-tests. Sample sizes: a, n=9; b, n=10; c, n=11.

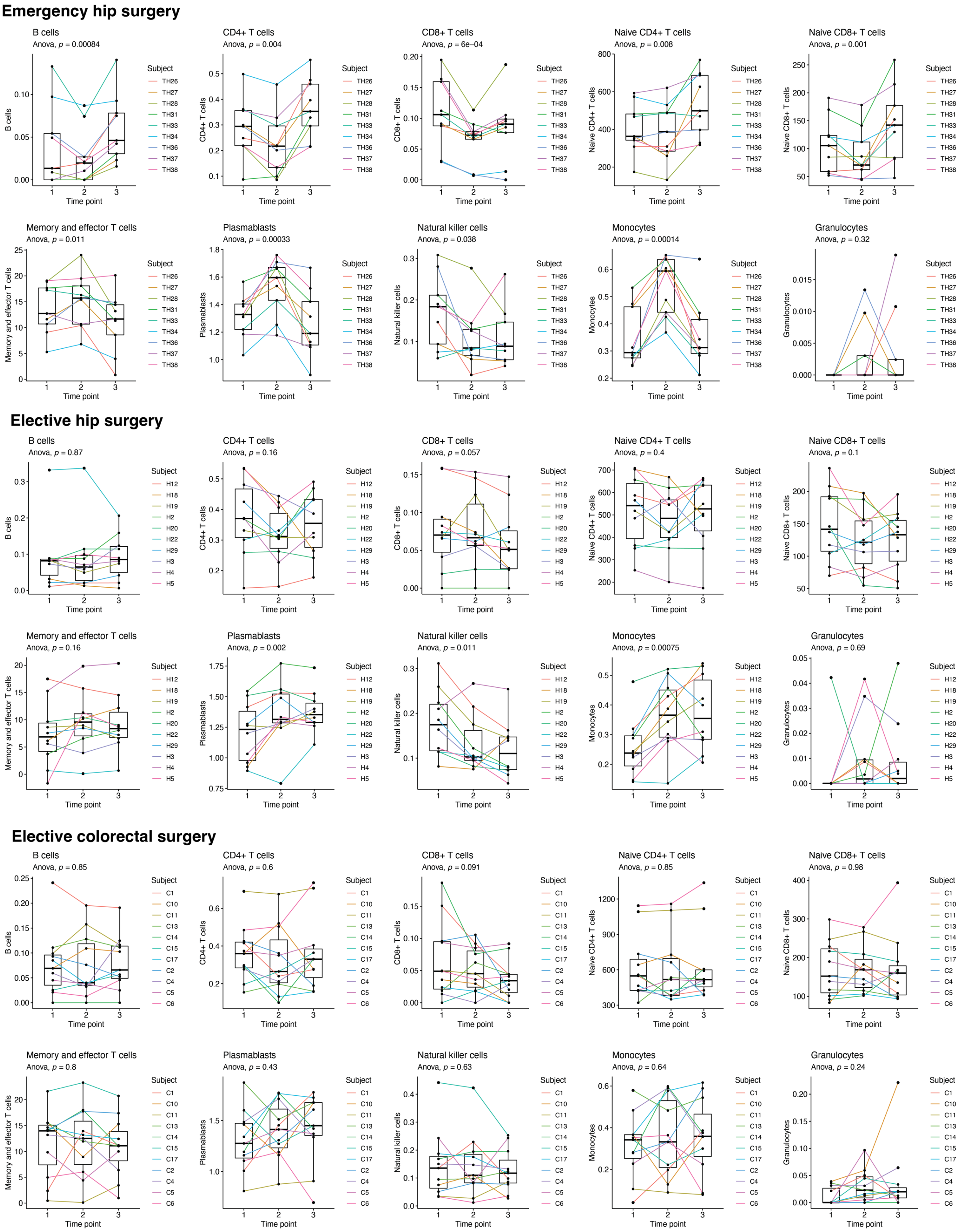

**Supplementary Figure 4. Predicted blood cell counts based on methylation data of patients undergoing major surgery.** Houseman[11] and Horvath/Levine[12] predictors of the indicated blood cell types were used to analyze blood composition dynamics in patients undergoing the indicated surgeries. P values were calculated with repeated-measures ANOVA. Sample sizes: emergency hip surgery, n=9; elective hip surgery n=10; elective colorectal surgery, n=11.

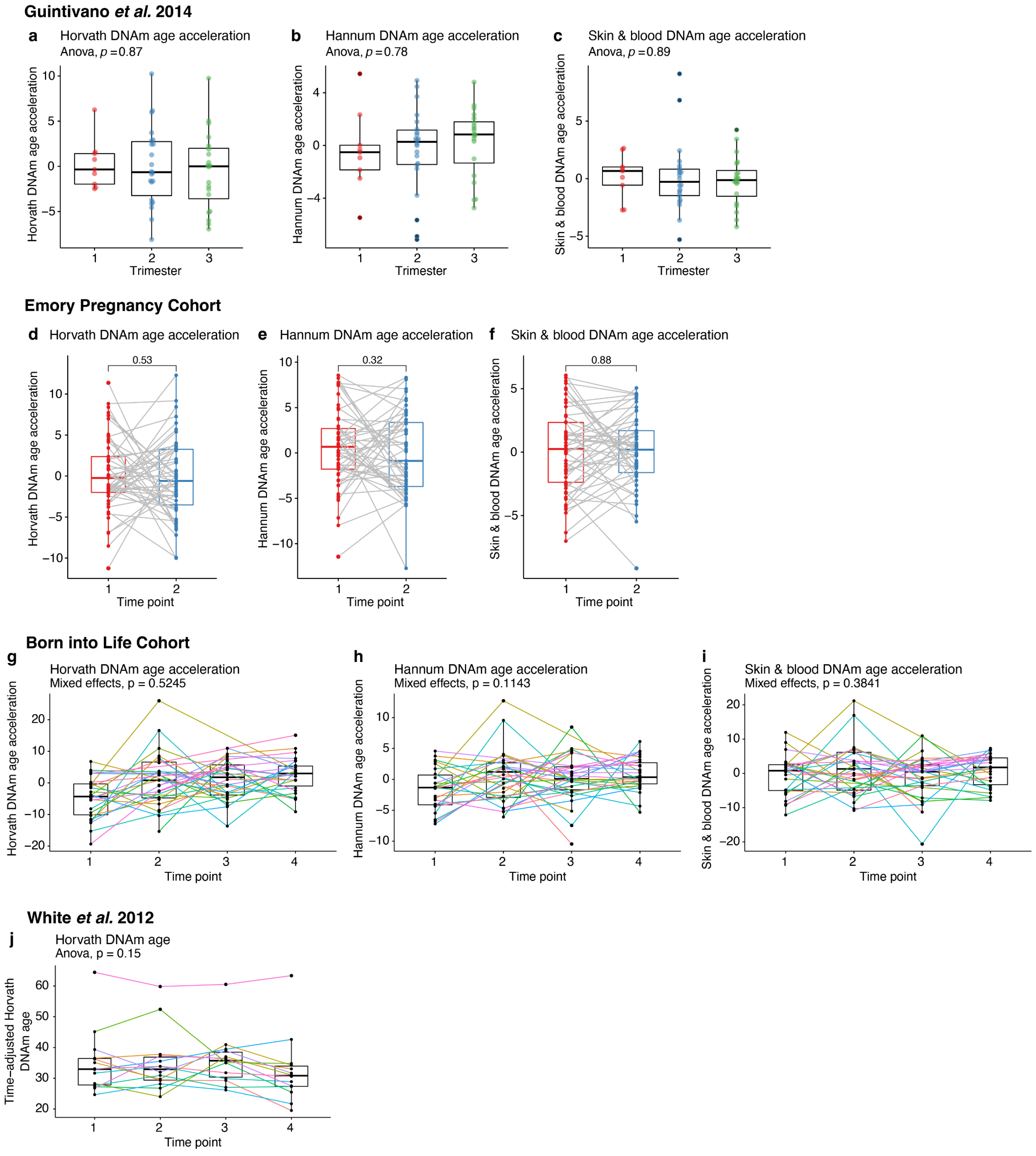

**Supplementary Figure 5. Horvath, Hannum, and Skin & Blood DNAm age biomarkers for human pregnancy datasets.** (a–c) Cross-sectional DNAm age acceleration analysis of pregnant Americans across the three trimesters of pregnancy using Horvath DNAm age (a), Hannum DNAm age (b), and Skin & Blood DNAm age (c). (d–f) DNAm age biomarkers (as in a–c) for a longitudinal study of pregnant African Americans with two blood samples collected over the course of pregnancy. Time point 1 corresponds to 7–15 weeks of pregnancy; time point 2 corresponds to 24–32 weeks of pregnancy. (g–i) DNAm age biomarkers (as in a–c) for Swedish mothers longitudinally tracked over the course of pregnancy. Time point 1 corresponds to pre-pregnancy; time point 2 corresponds to 10–14 weeks of pregnancy; time point 3 corresponds to 26–28 weeks of pregnancy; time point 4 corresponds to 2–4 days postpartum. (j) Horvath DNAm age (adjusted for the passage of time; see Methods for details) for a cohort of American mothers longitudinally tracked over the course of pregnancy and postpartum. Time point 1 corresponds to early pregnancy; time point 2 corresponds to mid-pregnancy; time point 3 corresponds to delivery; time point 4 corresponds to 6 weeks postpartum. P values were calculated using either repeated-measures ANOVA and paired t-tests or a mixed effects model with post-hoc pairwise comparison testing (see Methods). Sample sizes: a–c, n=9, 22, and 20 for trimesters 1, 2, and 3, respectively; d–f, n=53; g–i, n=33 total subjects who each provided up to 4 samples; j, n=14.

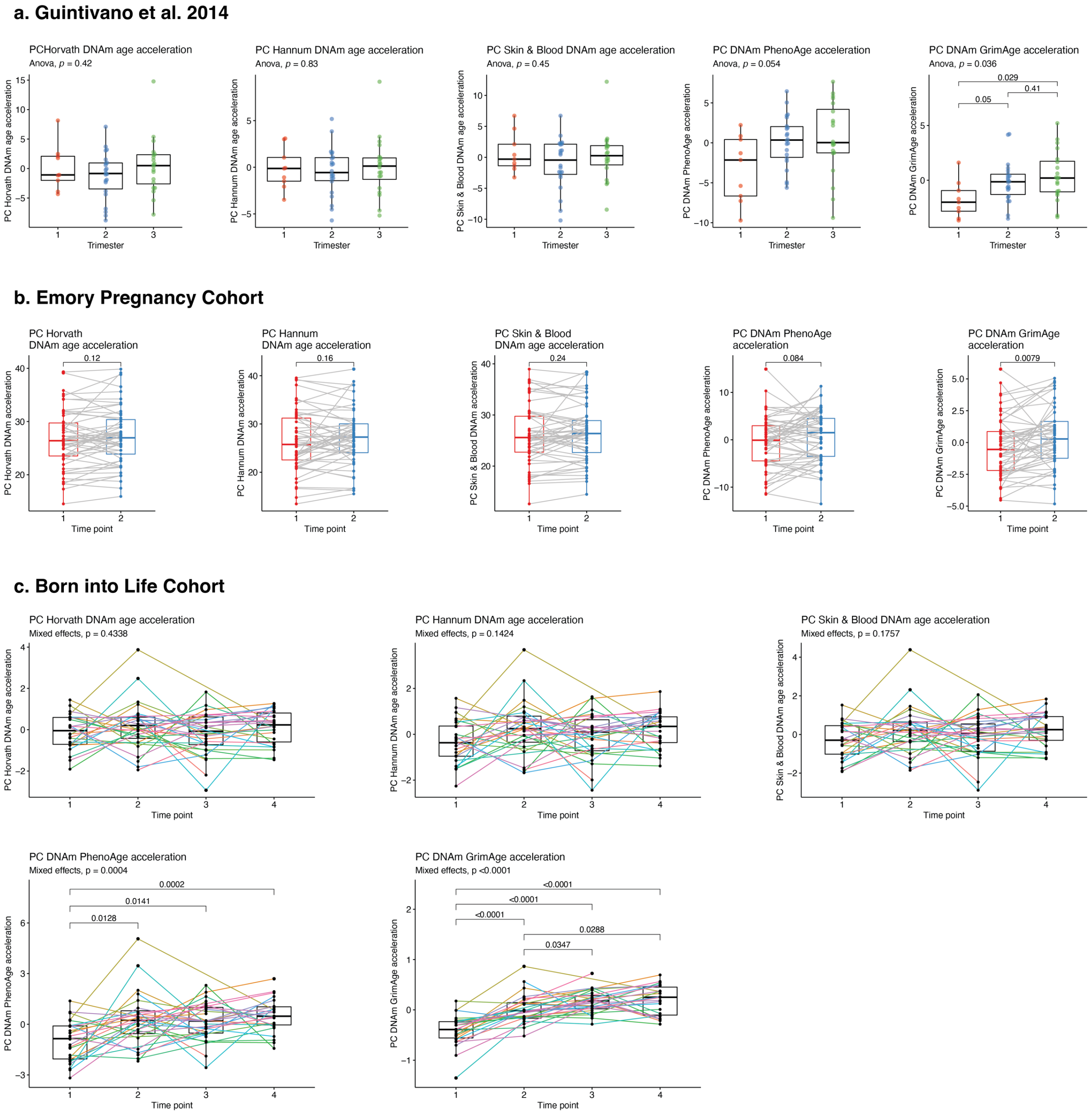

**Supplementary Figure 6. Principal component (PC) clocks results for human pregnancy cohorts.** PC clocks (Horvath DNAm age, Hannum DNAm age, Skin & Blood DNAm age, DNAm PhenoAge, and DNAm GrimAge) [10] were applied to methylation data from human pregnancy cohorts. P values were calculated with unpaired t-tests (a), paired t-tests (b), or mixed effects models (c). Sample sizes: a, n=9, 22, and 20 for trimesters 1, 2, and 3, respectively; b, n=53; c, n=33 total subjects who each provided up to 4 samples.

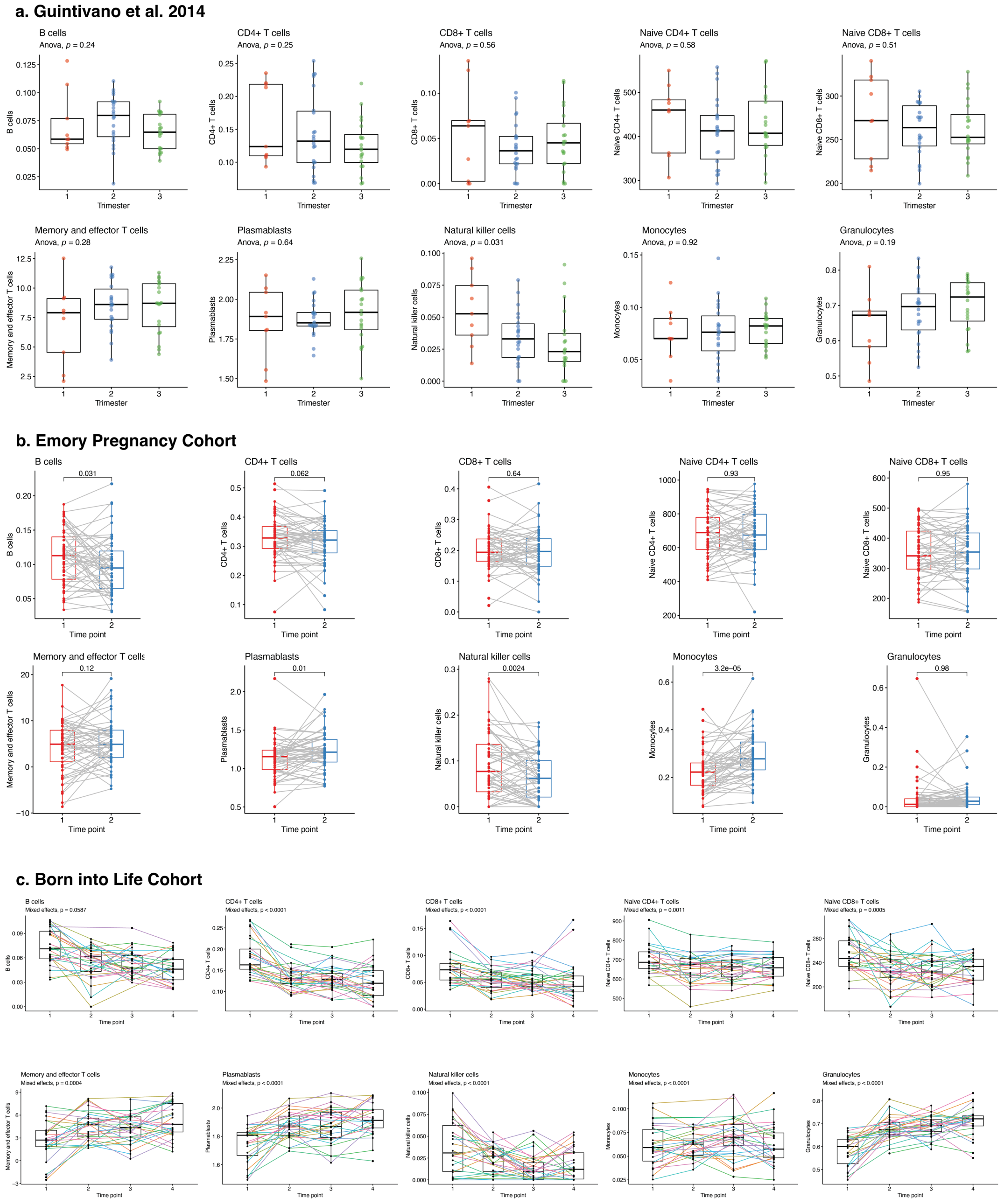

**Supplementary Figure 7. Predicted blood cell counts based on methylation data of human pregnancy cohorts.** Houseman[11] and Horvath/Levine[12] predictors of the indicated blood cell types were used to analyze blood composition dynamics during human pregnancy. P values were calculated with unpaired t-tests (a), paired t-tests (b), or mixed effects models (c). Sample sizes: a, n=9, 22, and 20 for trimesters 1, 2, and 3, respectively; b, n=53; c, n=33 total subjects who each provided up to 4 samples.

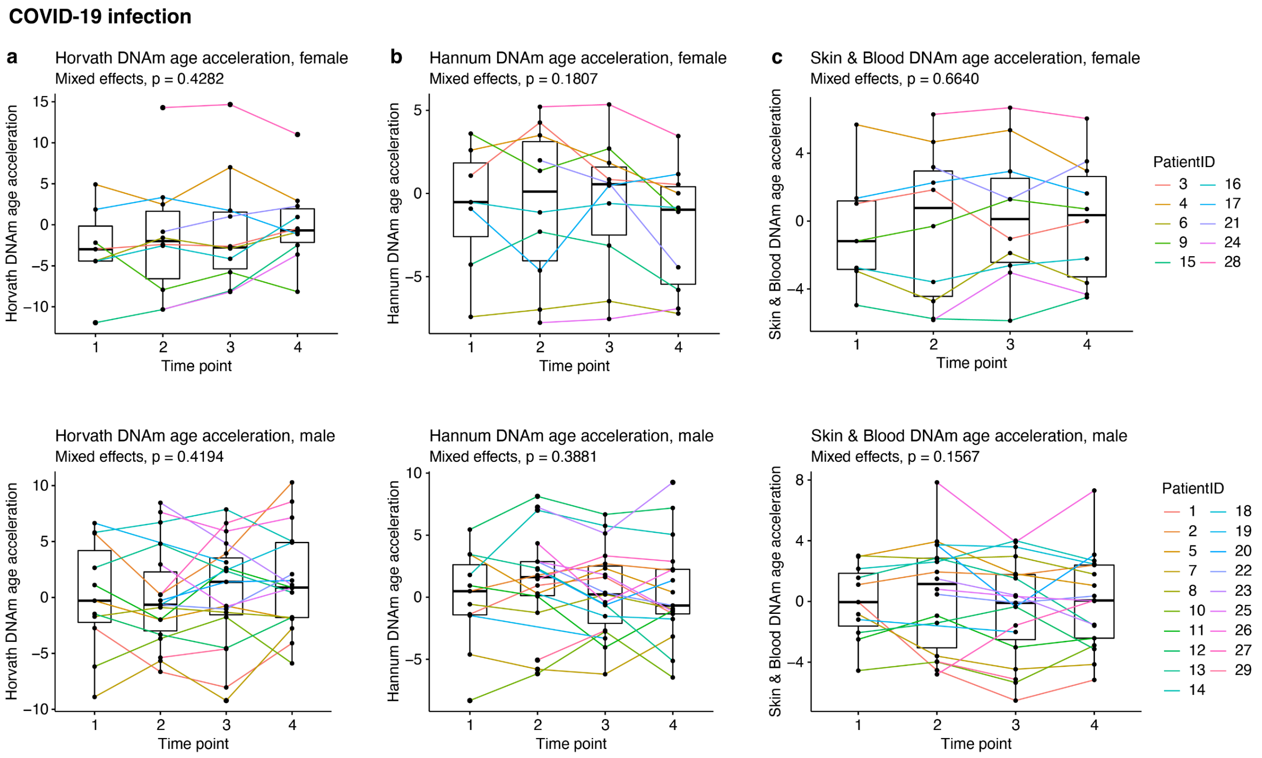

**Supplementary Figure 8. Horvath, Hannum, and Skin & Blood DNAm age acceleration for patients with COVID-19.** (a–c) DNAm age acceleration results for Horvath DNAm age (a), Hannum DNAm age (b), and Skin & Blood DNAm age (c). All upper panels show data for female patients and all lower panels show data for male patients. Time point 1 is within 5 days ICU admission; time point 2 is within 5 days of the midpoint of the ICU stay; time point 3 is within 5 days of the date of discharge from the ICU; timepoint 4 is ≥7 days post-ICU discharge. P values were calculated using a mixed effects model with post-hoc pairwise comparison testing. Sample sizes: a–c, n=10 female and n=19 male subjects total who each provided up to 4 samples.

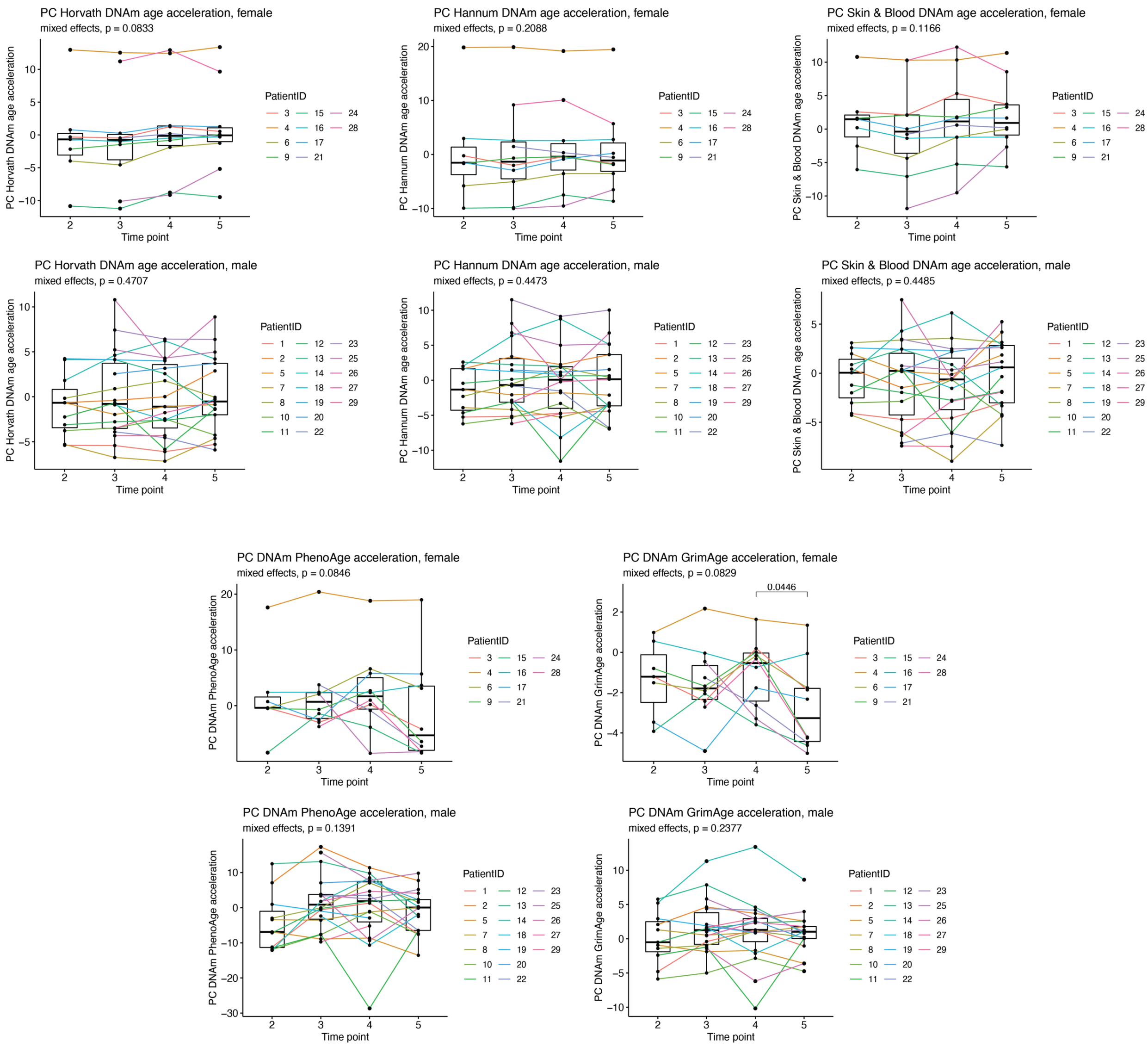

**Supplementary Figure 9. Principal component (PC) clocks results for COVID-19 patients.** PC clocks (Horvath DNAm age, Hannum DNAm age, Skin & Blood DNAm age, DNAm PhenoAge, and DNAm GrimAge) [10] were applied to methylation data from COVID-19 patients. P values were calculated with mixed effects models with post-hoc pairwise comparison testing. Sample sizes: n=10 female and n=19 male subjects total who each provided up to 4 samples.

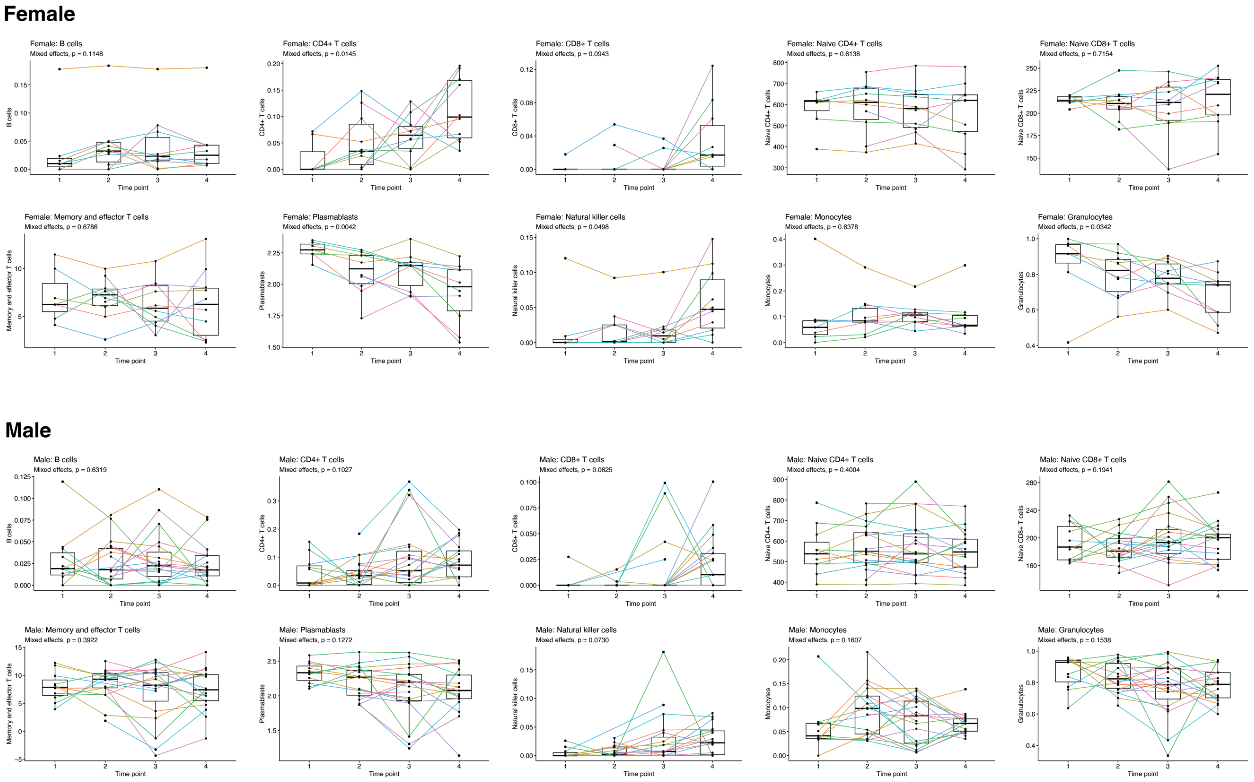

**Supplementary Figure 10. Predicted blood cell counts based on methylation data of COVID-19 patients.** Houseman[11] and Horvath/Levine[12] predictors of the indicated blood cell types were used to analyze blood composition dynamics during COVID-19 disease. P values were calculated with mixed effects models. Sample sizes: n=10 female and n=19 male subjects total who each provided up to 4 samples.

**Supplementary Table**

**Supplementary Table 1. Sources of human DNA methylation data.**

| **Dataset** | **Source** | **Description** | **Description of samples/time points:** | **Methylation array** |
|---|---|---|---|---|
| Sadahiro *et al.*[13] | GEO: GSE142536 | Longitudinal study of elderly adults undergoing major surgery | 1. Before surgery<br>2. Morning after surgery<br>3. Day of discharge from hospital (4–7 days post-surgery) | Illumina HumanMethylation450 BeadChip |
| Guintivano *et al.*[14] | GEO: GSE44132 | Cross-sectional study of pregnant women | Cross-sectional design with one sample per participant taken during a particular trimester | Illumina HumanMethylation450 BeadChip |
| Emory pregnancy cohort[15] | GEO: GSE107459 | Longitudinal study of pregnant women | 1. 7–15 weeks of pregnancy<br>2. 24–32 weeks of pregnancy | Illumina HumanMethylation450 BeadChip |
| Born into Life cohort[16,17] | Prof. Catarina Almqvist, Karolinska Institutet, Sweden | Longitudinal study of pregnant women | 1. Pre-pregnancy<br>2. 10–14 weeks of pregnancy<br>3. 26–28 weeks of pregnancy<br>4. 2–4 days postpartum | Illumina MethylationEPIC BeadChip |
| White *et al.*[18] | GEO: GSE37722 | Longitudinal study of pregnant women | 1. Early pregnancy<br>2. Mid-pregnancy<br>3. Delivery<br>4. 6 weeks postpartum | Illumina HumanMethylation27 BeadChip |
| COVID-19 | This study | Longitudinal study of patients undergoing intensive care for COVID-19 | 1. Within 5 days of admission to ICU<br>2. ±5 days of midpoint of ICU stay<br>3. Within 5 days of discharge from ICU<br>4. ≥7 days post-ICU discharge | Illumina MethylationEPIC BeadChip |
